# Selective degradation of ARF monomers controls auxin response in Marchantia

**DOI:** 10.1101/2022.11.04.515187

**Authors:** Shubhajit Das, Martijn de Roij, Simon Bellows, Wouter Kohlen, Etienne Farcot, Dolf Weijers, Jan Willem Borst

## Abstract

The plant signaling molecule auxin controls a variety of growth and developmental processes in land plants. Auxin regulates gene expression through a nuclear auxin signaling pathway (NAP) consisting of a ubiquitin ligase auxin receptor TIR1/AFB, its Aux/IAA degradation substrate, and the DNA-binding ARF transcription factors. While extensive qualitative understanding of the pathway and its interactions has been obtained by studying the flowering plant *Arabidopsis thaliana*, it is so far unknown how these translate to quantitative system behaviour *in vivo*, a problem that is confounded by large NAP gene families in this species. Here we used the minimal NAP of the liverwort *Marchantia polymorpha* to quantitatively map NAP protein accumulation and dynamics *in vivo* through the use of knock-in fluorescent fusion proteins. Beyond revealing the native accumulation profile of the entire NAP protein network, we discovered that the two central ARFs MpARF1 and MpARF2 are proteasomally degraded. This degradation serves two functions: it tunes the stoichiometry of auxin-responsive, positively acting MpARF1 and auxin-independent, negatively acting MpARF2, thereby permitting auxin response. Secondly, through mapping a minimal degradation motif, we found that degradation is likely selective for MpARF2 monomers and favours accumulation of dimers. Interfering with MpARF1:MpARF2 stoichiometry or preventing degradation of MpARF2 monomers caused strong growth defects associated with auxin response defects. Thus, quantitative analysis of the entire Marchantia NAP, allowed to identify a novel regulatory mechanism in auxin response, built on regulated ARF degradation.

## Introduction

The plant signaling molecule auxin triggers a multitude of growth, developmental and physiological responses across land plants^1,2^. Key to the cellular response is a nuclear auxin signaling pathway (NAP)^3^. Auxin promotes binding of AUXIN/INDOLE-3-ACETIC ACID (Aux/IAA) repressor proteins to the nuclear auxin receptor TRANSPORT INHIBITOR RESPONSE1/AUXIN SIGNALLING F-BOX (TIR1/AFB)^4^ This triggers degradation of Aux/IAA proteins^5^ and releases DNA-binding AUXIN RESPONSE FACTORS (ARFs)^6,7^ from inhibition (Figure 1.A). In the past decades, the nuclear auxin pathway has been studied extensively in the angiosperm *Arabidopsis thaliana*^8^. These studies have led to a comprehensive qualitative model of auxin signaling, supported by atomic structures of each component. A key challenge in understanding auxin response is the large size of gene families representing each component in angiosperms^9^. Specifically, it is hard to tell if any one protein behaves typically or atypically. More recent analysis of the auxin response system in the bryophytes *Physcomitrium patens* (a moss) and *Marchantia polymorpha* (a liverwort) has helped reduce system complexity and derive common, core principles. Phylogenetic analysis showed that the simplest NAP systems consist of one ARF in each of the three subclasses (A, B and C), a single Aux/IAA and a single receptor, in addition to a non-canonical ARF member^10^. Studies in Physcomitrium and Marchantia led to the model that auxin response revolves around antagonistic interactions between A- and B-class ARFs, competing for the same DNA binding sites. While A-ARFs are regulated by auxin and can switch between repression and activation, B-ARFs are auxin-independent repressors^11,12^. Thus, stoichiometry of the A and B-class ARFs is predicted to determine the output of auxin response.

**Figure 1:**
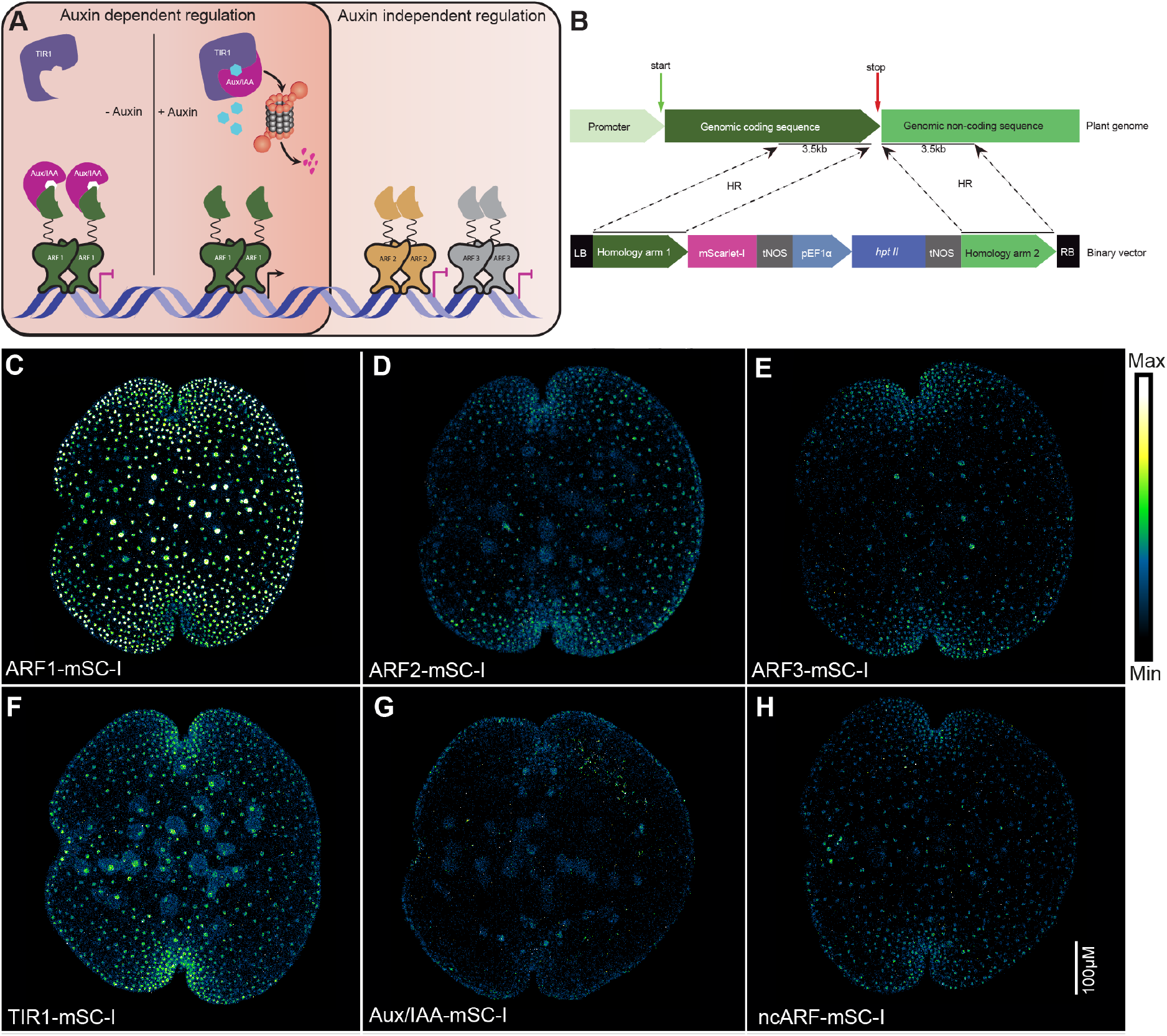
(A) Schematic diagram of nuclear auxin signaling pathway. In absence of auxin, Aux/IAA (AUXIN/INDOLE-3-ACETICACID) repressors interact with ARFs (AUXIN RESPONSE FACTOR) to inhibit transcriptional activation. In presence of auxin, TIR1(TRANSPORT INHIBITOR RESPONSE1) interacts with Aux/IAA and targets it for proteasomal degradation. Free ARFs initiate gene transcription of auxin response genes. ARF2 is independent from this regulation. Regulation mode for ARF3 is not well characterized. Both ARF2 and ARF3 acts as transcriptional repressors. (B) Principle of homologous recombination mediated genomic knock-in of fluorescent proteins at the C-terminus of a gene of interest. (C-H) Native expression patterns of core nuclear auxin signaling proteins in dormant gemmae of Marchantia.

With all the deep knowledge about the NAP, one key element is largely unresolved: essentially all *in vivo* findings are built on qualitative data, and the true *in vivo* concentrations and accumulation patterns of NAP components, and their relative concentrations or stoichiometries are unknown. Given that any biochemical interaction is determined both by affinity and concentration, it will be essential to map protein patterns *in vivo*. The facile homologous recombination-based gene targeting in bryophytes allows engineering lines in which NAP components are e.g., fluorescently labelled *in situ*.

In this study, we generated fluorescent genomic knock-in lines for all NAP proteins. Beyond resolving spatial and temporal maps of protein accumulation, this strategy allowed us to identify active proteosomal degradation of both MpARF1 (A-class) and MpARF2 (B-class). We show that this degradation actively tunes A/B ARF stoichiometry to allow normal development and auxin response. By mapping the degron, we discovered that monomeric MpARF2 is selectively degraded, and that this is required for normal development. Our study provides a resource for quantitative analysis of auxin response in Marchantia and reveals novel modes of regulating ARF activity through selective degradation.

## Results

### Development of a collection of genomic knock-in lines of the Marchantia NAP

Many activities and properties underlying *in vivo* protein function are strongly concentration dependent. Therefore, quantitative understanding of any biological process requires monitoring endogenous proteins accumulation patterns. To quantify the native accumulation patterns of the entire auxin response system in a land plant, we generated genomic knock-in translational fluorescent fusion lines in *Marchantia polymorpha* (Figure 1.B)^13^. Using homologous recombination, we knocked-in a fluorescent protein (either mNeonGreen or mScarlet-I) at the C-terminus of all auxin signaling proteins: MpARF1 (class A), MpARF2 (class B), MpARF3 (class C), MpAux/IAA, MpTIR1 and MpncARF. Either overexpression or loss of function for all these proteins causes strong phenotypes^10,11,14^. All knock-in lines displayed wild-type-like morphologies and responded to externally applied auxin like wild-type (Supplementary Figure 1). This suggests that all fusion proteins are fully functional, and that none accumulates at levels outside of the normal range. Next, we used confocal microscopy to visualize all auxin signaling proteins in dormant gemmae residing within the gemma cup and compared fusions of all NAP components to the same fluorescent protein (mScarlet-I) under identical conditions. The three ARFs showed clear differences in both their qualitative and quantitative accumulation patterns, and all were localized to nuclei, as predicted (Figure 1.C-E). MpARF1-mScarlet-I accumulated most strongly among all NAP components and was present in most cell types of the gemma. MpARF2-mScarlet-I accumulation was much weaker and was only expressed in the apical and green cells but not in region distal from the apical notch where rhizoids and air chambers reside. MpARF3-mScarlet-I (class C) displayed a weak yet broad accumulation pattern in all cell types. The MpTIR1-mScarlet-I receptor and the non-canonical MpncARF-mScarlet-I showed low, but nuclear accumulation, while MpAux/IAA-mScarlet-I could not be detected (Figure 1.F-H). Thus, we established a collection of knock-in lines that allow to derive *in vivo* NAP protein accumulation patterns.

### Dynamics of NAP protein accumulation

MpAux/IAA-mScarlet-I was undetectable at dormant stage gemma. We therefore checked protein accumulation in the first 24 hours following germination using a microscope slide mount to simultaneously grow and image the same gemma. (Supplementary Figure 2). We did not observe any fluorescence signal in the first 8 hours but detected a weak nuclear signal after 24 hours (Figure 2A). It is conceivable that the MpAux/IAA protein is continuously degraded due to high auxin concentrations at earlier stages. Indeed, treating dormant gemmae with the proteasome inhibitor MG132 led to a clear nuclear signal within 2 hours (Figure 2B). We measured free IAA concentrations in germinating gemma, and found that these indeed start high, and drop off as the gemma grows (Figure 2D), consistent with IAA-triggered degradation at early stages. To determine causality, we blocked auxin synthesis combining L-Kynurenine and Yucasin inhibitors^15,16^. Nuclear MpAux/IAA-mScarlet-I accumulation was detected across the gemma within 2 hours of treatment (Figure 2C). These findings suggest that MpTIR1 is active throughout gemma development in mediating IAA-triggered MpAux/IAA degradation. Indeed, MpTIR1-mScarlet-I was detectable at all stages but showed a gradual rise in protein levels as gemmae grew (Figure 2E). Thus, as gemmae break dormancy and grow, the auxin response system moves from restrictive to permissive.

**Figure 2:**
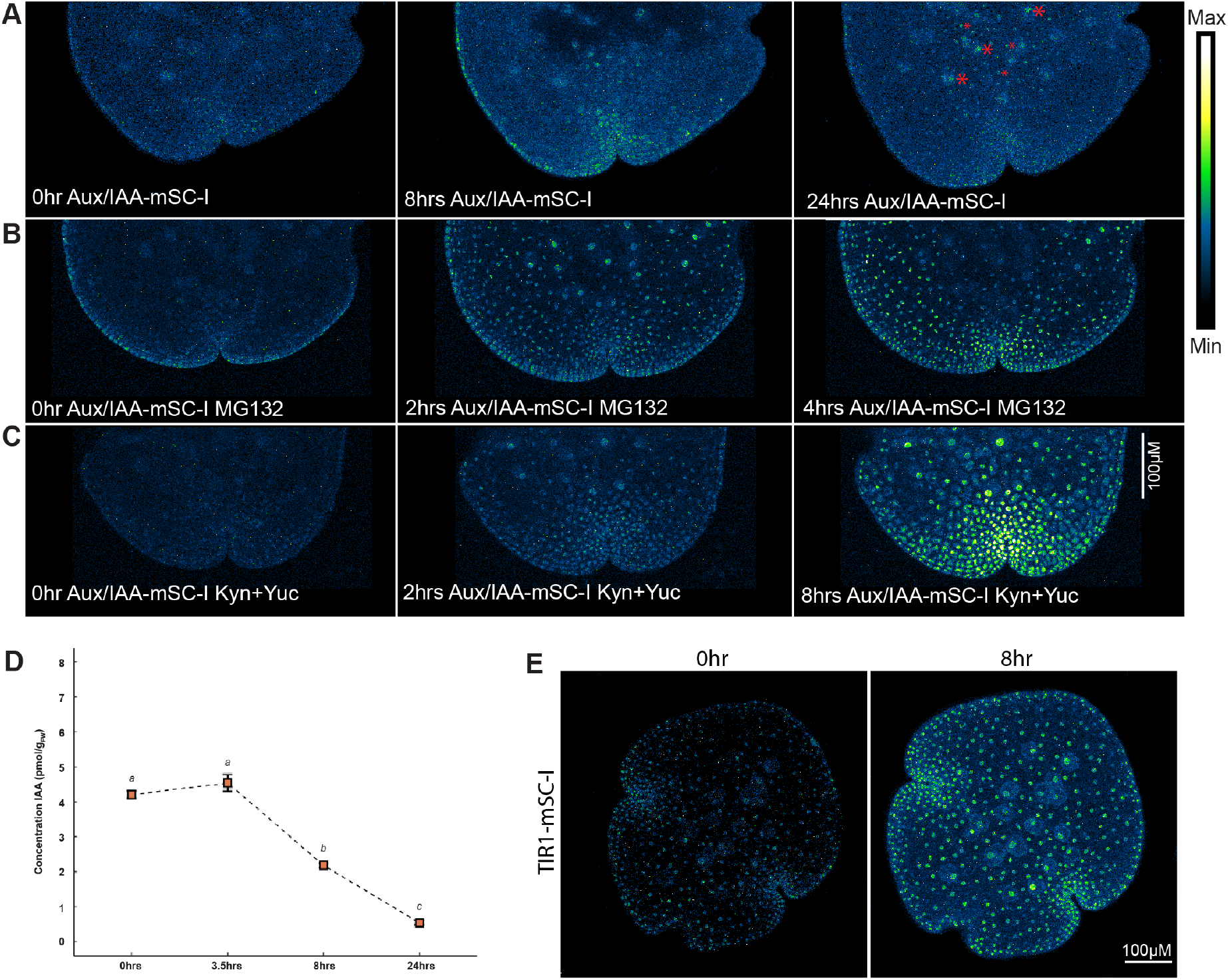
(A)Time course imaging of Aux/IAA-mSC-I with mock, (B)100μM MG132, and (C) 50μM Kynurenine and 50μM Yucasin. Low expression of Aux/IAA at 24 hours in mock treated sample is highlighted with red asterisks in (A). (D) Concentration dynamics of free IAA in gemmae during dormancy to germination. LC-MS quantification shows total IAA levels drop with gemma germination. (E) Time course imaging of TIR1-mSC-I shows that the expression levels gradually increase after gemma germination.

To explore the temporal dynamics of MpARF expression, we imaged each ARF during the first 24 hours following germination. Remarkably, while MpARF3 patterns did not change in this time window, we found both MpARF1 and MpARF2 protein levels to progressively decline after germination (Supplementary Figure 3.A-C). Given the higher starting levels of MpARF1 in dormant gemmae, and lower MpARF2 concentrations, the former was still detectable after 8 hours, while MpARF2 signal declined to undetectable levels (Supplementary Figure 3.A-B). Comparable dynamics were found for the MpARF1-mNeonGreen and MpARF2-mNeonGreen proteins (Supplementary Figure 3.D), indicating that behavior is independent of the protein tag. Expressing unfused nuclear mScarlet-I or mNeonGreen from a constitutive EF1α promoter (pEF1α∷NLS-mScarlet-I; pEF1α∷NLS-mNeonGreen) led to stable accumulation patterns (Supplementary Figure 3.E) showing that this phenomenon is specific to MpARF1 and MpARF2.

### Regulated degradation controls ARF stoichiometry for normal development and auxin response

To test whether ARF level decline was transcriptionally regulated, we quantified ARF transcripts in germinating gemmae. Neither of the MpARF transcripts changed significantly over the first 8 hours after germination (Figure 3.A), suggesting that MpARF1/2 protein levels are post-transcriptionally controlled by protein-level regulation. Indeed, both proteins accumulated to much higher levels when treated with the proteasome inhibitors MG132 (Figure 3.B-C, E) or Bortezomib (Supplementary Figure 4), while MpARF3 was unaffected by the treatments (Figure 3.D-E). Importantly, proteasome inhibition completely prevented the decline of MpARF1 and MpARF 2 signal (Figure 3.B-C, E). This implies that both MpARF1 and MpARF2, but not MpARF3 are proteasomally degraded.

**Figure 3:**
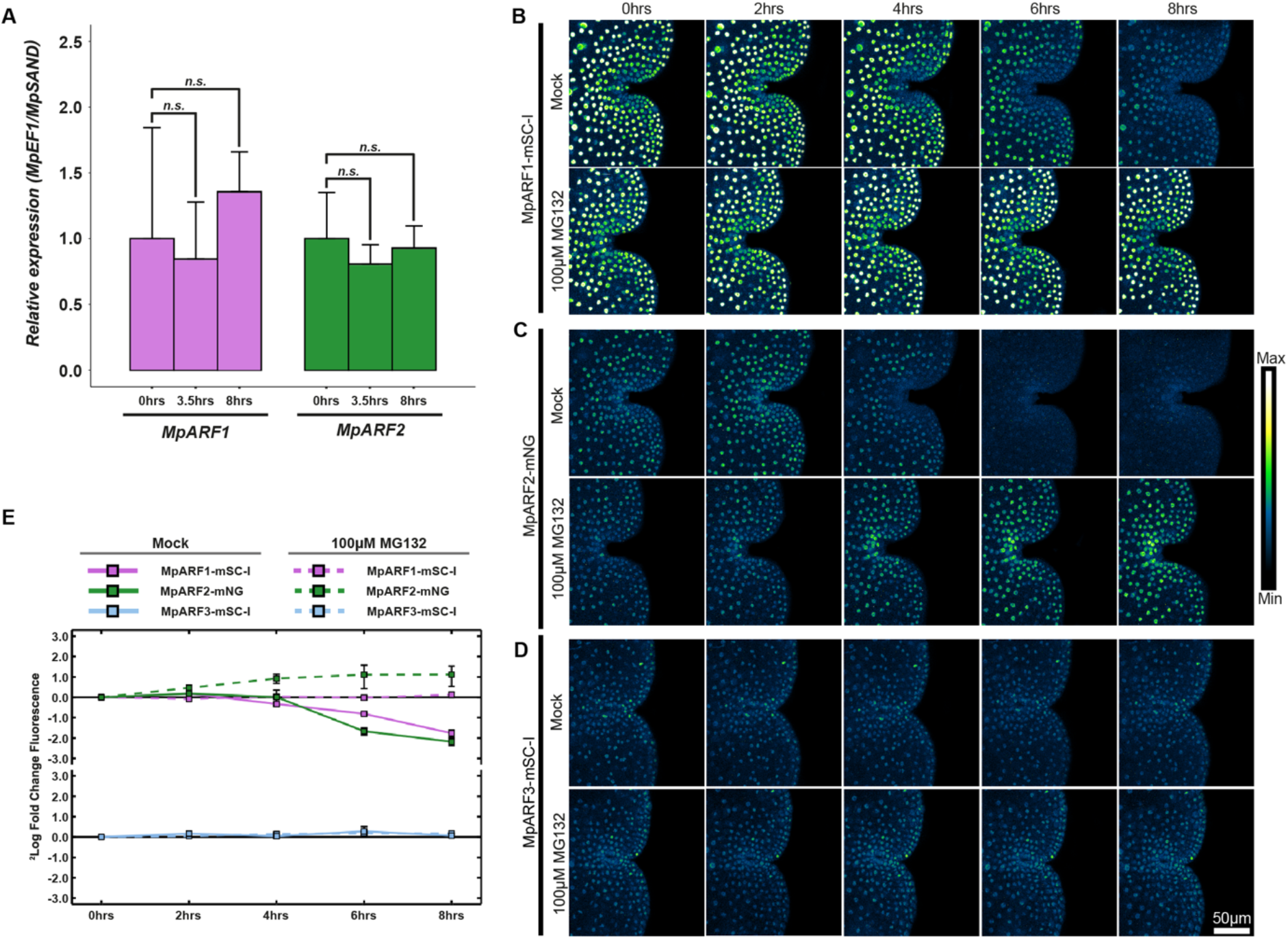
(A) qRT-PCR of MpARF1 and MpARF2 shows stable gene transcription during gemma germination. (B-D) Time-course imaging of ARF1, ARF2 and ARF3 in germinating gemmae. Both ARF1 and ARF2 fluorescence signal starts to decline after germination. ARF2-mNG fluorescence is undetectable after 6 hours, whereas ARF1-mSC-I expression is maintained at 8 hours. Upon treatment with proteasome inhibitor 100μM MG132, the signal decline of ARF1-mSC-I and ARF2-mNG is blocked, implying proteasomal regulation of ARF1 and ARF2. ARF3-mSC-I expression remains stable in both mock and 100μM MG132 treated samples, suggesting that ARF3 lacks proteasomal regulation unlike ARF1 and ARF2. (E) Quantification of ARF1, ARF2, and ARF3 protein expression fold changes in mock and 100μM MG132 treated samples.

A key question is what function this regulated ARF degradation serves. Given that we previously proposed that auxin responsiveness in Marchantia is determined by the stoichiometry of MpARF1 and MpARF2^11^, one option would be that this stoichiometry is actively regulated. Since such stoichiometries may differ per cell, it is not possible to derive them from comparing individual knock-in lines. We therefore crossed MpARF1-mScarlet-I with MpARF2-mNeonGreen and MpARF1-mNeonGreen with MpARF2-mScarlet-I and obtained two double knock-in lines. The accumulation profiles of both MpARFs in whole gemma of these double knock-ins were the same as observed for the single ARF knock-ins (Figure 4.A-B; Supplementary Figure 3.A-B, D). As predicted from their individual patterns, MpARF1 and MpARF2 are present in differing stoichiometries in different cell types. In meristematic apical notch cells, MpARF2 displayed higher accumulation than MpARF1. (Figure 4.C). In contrast, in rhizoid initial cells, MpARF2 protein was near-undetectable whereas MpARF1 showed clear accumulation (Figure 4.D). In the region separating apical notch and the rhizoid domain, we observed presence both MpARFs (Figure 4.B).

**Figure 4:**
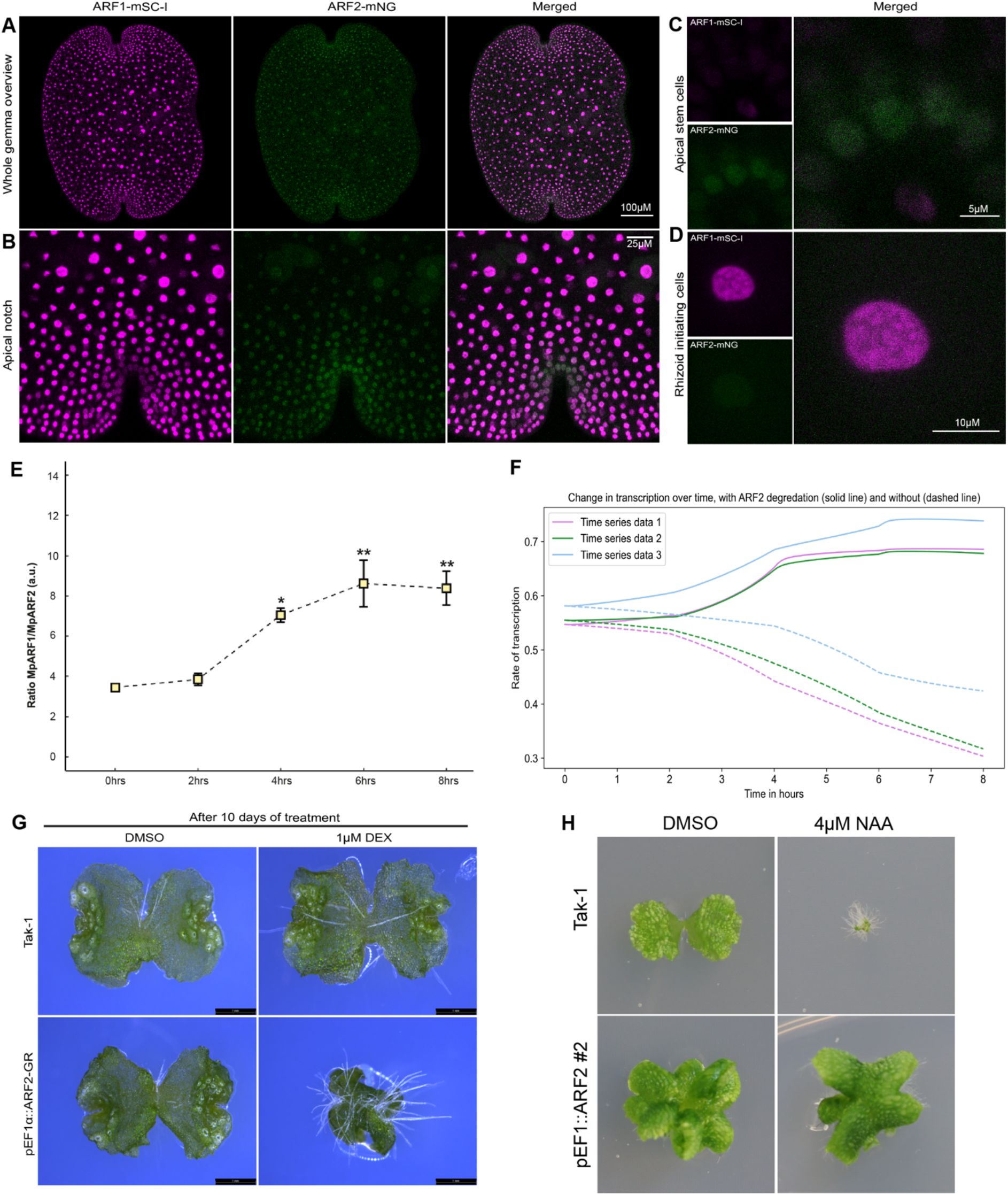
(A) Overview of ARF1-mSC-I and ARF2-mNG accumulation patterns in dormant gemmae of double knock-in lines. (B) ARF1-mSC-I and ARF2-mNG expression pattern in apical notch region in dormant gemma of double knock-in line. (C) ARF2-mNG expression in apical notch cells, ARF1-mSC-I expression is not detectable in these cells. (D) ARF1-mSC-I expression is high in rhizoid initial cells. ARF2-mNG expression if negligible in these cells. (E) Quantification of ARF1:ARF2 stoichiometry during gemma germination in ARF1-mNG and ARF2-mNG single knock-in lines. (F) Mathematical model predicted transcription pattern of an auxin inducible gene during gemma germination. Solid lines indicate transcription rate in normal condition, whereas dashed lines indicate transcription rate in absence of proteasomal degradation of ARF2. (G) Inducible ARF2 overexpression leads to retarded growth. pEF1∷ARF2-GR lines were treated with 1μM dexamethasone and plant growth was compared with wild-type. After 3 days of DMSO treatment, both wild-type Tak-1and pEF1∷ARF2-GR lines grew similarly. However, on dexamethasone treatment, induced ARF2-GR overexpression lines growth was affected negatively in comparison to wild-type. (H) Constitutive overexpression of ARF2 in pEF1∷ARF2 line leads to auxin resistance.

We next determined MpARF1/MpARF2 stoichiometries following gemmae germination and observed that the different MpARF starting concentrations and degradation rates result in a progressive increase of MpARF1:MpARF2 stoichiometry after gemmae germination (Figure 4.E), favouring MpARF1.

We next asked what the impact of this change in MpARF1:MpARF2 stoichiometry may be, and first developed a mathematical model of the minimal Marchantia NAP and its known interactions (Supplementary File 1). Using a piecewise polynomial function, we fitted quantified MpARF accumulation profiles into the mathematical model, where the effect of changing MpARF1 and MpARF2 levels on the outcome of auxin response, was modelled as the transcription of an ARF-regulated gene. Model simulations predicted that the MpARF1:MpARF2 ratio increase would enhance transcriptional response output (Figure 4.F). Conversely, simulated loss of MpARF2 degradation predicted an opposite effect on response output (Figure 4.F). Thus, MpARF2 degradation may be required to switch from an inactive to an active transcriptional state of auxin-regulated genes.

To test the prediction that regulated, low MpARF2 levels are needed for activating auxin response in gemmae, we either stably or inducibly overexpressed MpARF2 in wild-type background. Inducing MpARF2-GR activity with Dexamethasone prevented thallus growth (Figure 4.G) and formation of rhizoids (Supplementary Figure 5), a reliable marker for auxin response output^17,18^. Likewise, 2-week-old stable pEF1-MpARF2 overexpression lines had multiple apical notches, were strongly defective in their growth, lacked gemma cups (Supplementary Figure 6) and were completely insensitive to auxin treatment (1-Napthaleneacetic acid; 1-NAA; Figure 4.H; Supplementary Figure 6). These results support the prediction that elevated levels of MpARF2 inhibit plant growth and auxin response and therefore targeted removal of ARF2 is required to initiate growth.

### Identification of a degron driving ARF degradation

Both MpARF1 and MpARF2 are rapidly degraded following germination, and MpARF2 hyper-accumulation by overexpression suggests that developmental progression is very sensitive to MpARF2 levels. As a first step in dissecting mechanisms underlying regulated degradation, we therefore focused on identifying requirements for MpARF2 degradation. We first expressed each of the three conserved MpARF2 domains as an mNeonGreen fusion from an MpARF2 promoter fragment and added a nuclear localization signal to ensure nuclear entry (Figure 5.A). As a control we included full-length MpARF2 and found it to behave as in knock-in lines. Both the Middle Region (MR) or the Phox-Bem1 (PB1) strongly accumulated at both 0 hours and 8 hours after germination (Figure 5.A), suggesting that these do not carry a degradation signal. In contrast, almost no accumulation was detected for the DNA-Binding Domain (DBD), either at dormant stage or after 24 hours (Figure 5.A). However, MG132 treatment caused protein to accumulate (Figure 5.B), confirming that the DBD carries a signal that is sufficient for MpARF2 degradation. Interestingly, the DBD alone was much less stable at dormant stage than full-length protein (Figure 5.A), suggesting that regions outside the DBD protected MpARF2 from degradation.

**Figure 5:**
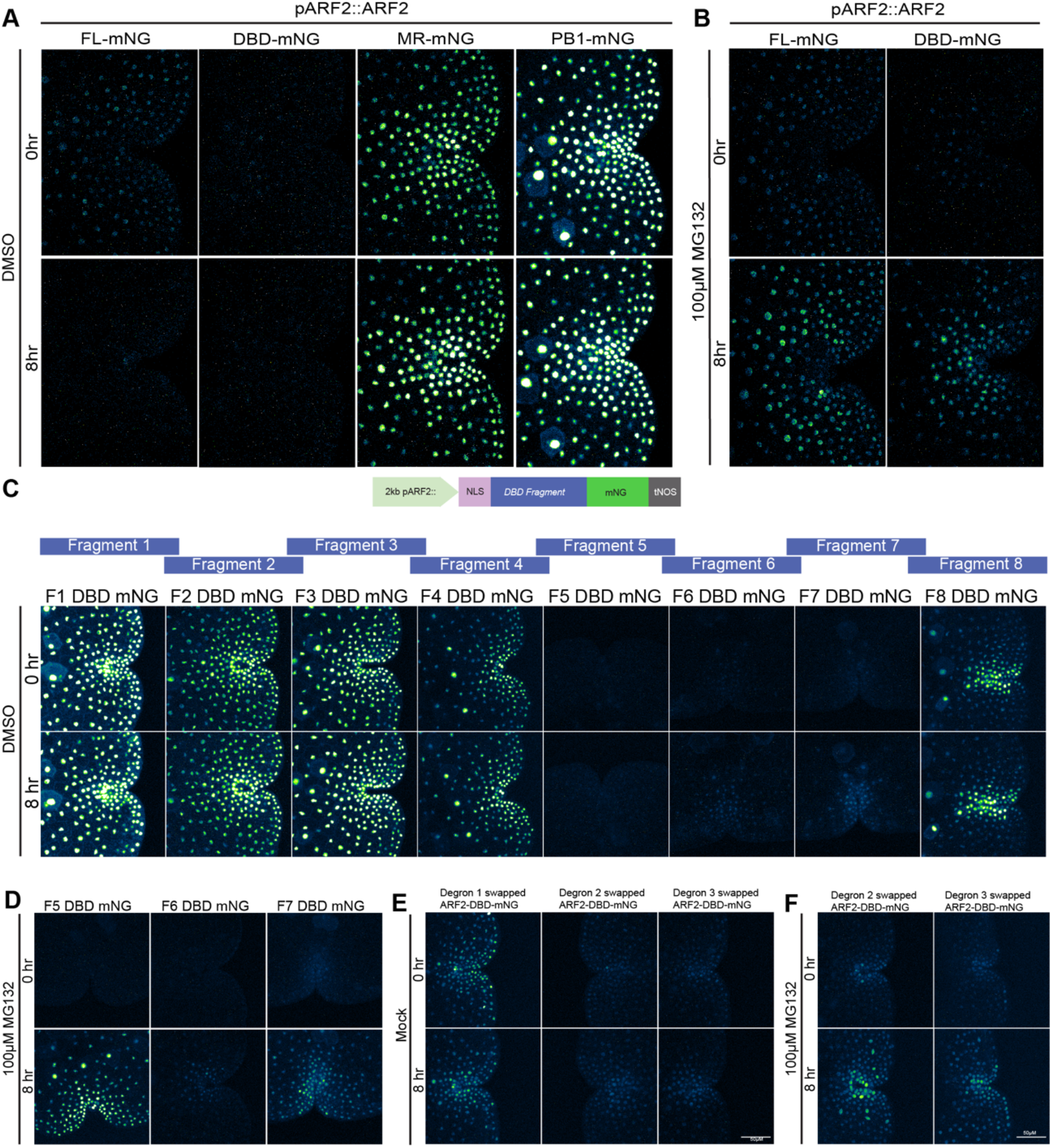
(A) ARF2 domain split analysis. Each domain of ARF2 fused to mNG was expressed separately to determine the location of the degradation signal in the protein. Both the MR and the PB1 domain show high stability at 0 and 8 hours. The full-length ARF2 is stable at dormant stage, but it is degraded after germination. The DBD is degraded already at dormant stage. (B) 100μM MG132 treatment can inhibit degradation of both full-length and ARF2-DBD. (C) ARF2-DBD fragment analysis to narrow down the location of the degron. ARF2-DBD was divided into 8 overlapping fragments and fused to mNG. Expression of each fragment in plants shows that fragments 5,6 and 7 are instable in dormant stage, whereas all other fragments are stable after 8 hours. (D) 8 hours of 100μM MG132 treatment on fragments 5,6 and 7 is sufficient to block degradation of these fragments. (E) ARF2 DBD degron swap experiment. The 25 amino acid overlapping region between ARF2-DBD fragment 6 and 7 were divided into three small motifs. Each small motif was named as degron 1, 2 and 3. We swapped these small motifs in ARF2-DBD with homologous sequences from ARF3-DBD, which lacks proteasomal regulation. Swapping degron 2 and degron 3 with ARF3 sequence is not sufficient to stop ARF2-DBD degradation. Only when the degron 1 (the first 12 amino acids in the overlap between fragment 6 and 7) is replaced, ARF2-DBD becomes stable both at dormant stage and after 8 hours of growth. (F) MG132 treatment blocks the degradation of degron 2 and degron 3 swapped ARF2-DBD, confirming their proteasomal degradation.

To identify a minimal degron motif within the MpARF2 DBD, we tiled the DBD into 8 short fragments of 75 amino acids, carrying 25 overlapping residues with neighbouring fragments (Figure 5.C), and expressed each as we did the protein domains. All DBD fragments, except fragments 5, 6 and 7 showed strong and stable nuclear accumulation (Figure 5.C). Fragments 5, 6 and 7 did not show accumulation, but treatment with MG132 caused protein to accumulate (Figure 5.D). This identifies the region encompassed by these three fragments as a degron sufficient for MpARF2 degradation.

These experiments demonstrate sufficiency of these regions for degradation, but do not tell if the degron is required in the context of the larger domain. We therefore divided the 25 amino acid overlapping region between fragment 6 and 7 into 3 smaller overlapping motifs of 12 amino acids each and swapped each with the equivalent sequences from the non-degradable MpARF3 protein in the context of MpARF2-DBD-mNeonGreen. Among these, only the first swap led to stable accumulation of ARF2-DBD, while the other two were degraded (Figure 5.E-F). Thus, this delineation identifies a minimal region in the MpARF2 DBD that is both necessary and sufficient for regulated degradation.

### Selective degradation of ARF2 monomers is required for normal development

Previously, we solved the crystal structure of the MpARF2 DBD^11^, and we used this structure to map the location of the minimal fragments that can drive MpARF2 degradation. All three fragments mapped to the homodimerization interface (Figure 6.A). ARF DBD dimerization is required for high-affinity DNA binding in Arabidopsis^19,20^, and Arabidopsis ARF5/MP mutants that cannot dimerize through their DBD fail to complement the mutant^20^. The location of the degron at this interface prompts the hypothesis that MpARF2 degradation selectively targets monomers, since the degron would be occluded in the dimer. To test this hypothesis, we engineered a dimerization mutation into the MpARF2-DBD (ARF2^G286I^) and assessed stability. As predicted, this mutant is highly unstable, and cause loss of protein accumulation even in dormant gemmae (Figure 6.B). In addition to the DBD, most ARFs have a C-terminal PB1 domain that can also serve as homotypic dimerization domain. These two dimerization domains likely act cooperatively, with each favouring dimerization at the other domain. Given that the DBD alone is less stable than the full-length MpARF2, we explored if dimerization through the PB1 domain indirectly stabilizes MpARF2. We therefore engineered a PB1 dimerization mutant (ARF2^K760S, OPCA^), and found this to render MpARF2 highly unstable (Figure 6.B). Likewise, combining the ARF2^G286I^ and ARF2^K760S, OPCA^ mutations led to similar instability compared to each single mutant (Figure 6.B), yet MG132 treatment allowed mutant proteins to accumulate (Figure 6.C). This is consistent with cooperative dimerization preventing degradation of MpARF2.

**Figure 6:**
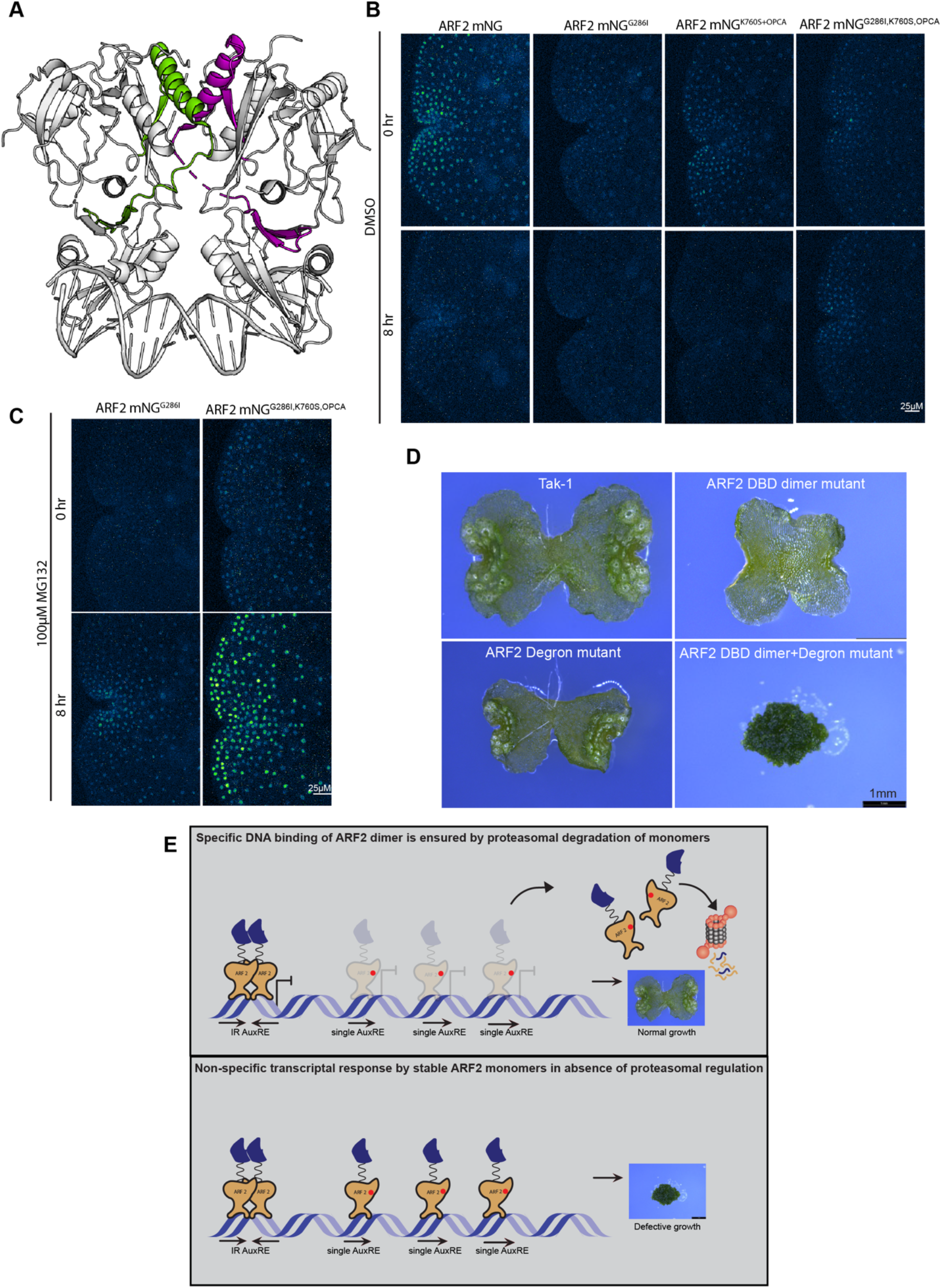
(A) Mapping the fragments 5,6 and 7 on the crystal structure of ARF2-DBD shows that the degron is in the DBD dimer interface. (B) Time-course imaging of ARF2 dimerization mutants. Disruption of DBD dimerization (ARF2^G286I^), PB1 dimerization (ARF2^K760S+OPCA^) and double dimerization (ARF2^G286I+K760S+OPCA^) in ARF2 causes degradation of the full-length proteins in dormant stage. (C) Treatment with proteasomal degradation inhibitor results in accumulation of both DBD dimerization mutant and DBD PB1 double dimerization mutant proteins. (D) Severe growth defects in plants expressing ARF2 with DBD dimer mutation and degron mutation (ARF2^G286I+F6F7 swapped^). The mutant does not have any differentiation of tissue and grows as a mass of undifferentiated callus. This shows the effect of a stable ARF2 monomer in plants and highlights the importance of ARF2 degradation by proteasome. (E) Proposed model showing the necessity of a monomer specific degradation mechanism for specific auxin response.

If the MpARF2 dimerization interface occludes the degron, and allows for selective degradation of monomeric MpARF2, a key question is what function this serves. We therefore engineered a version of MpARF2 in which both DBD dimerization and degradation are prevented without affecting any other property of the protein, and expressed it from an ARF2 promoter fragment. To prevent DBD dimerization, we used the G286I mutation, and the swap of sub-degron region 1 from MpARF3 was used to prevent degradation. This should lead to the stable accumulation of monomeric MpARF2. This MpARF2 double mutant showed a striking phenotype, lacking a flat thallus, rhizoids, gemmae or gemma cups, whereas the individual DBD dimerization mutant or degron region 1 swap showed mild effects on growth (Figure 6.D). The double mutant grew as a mass of undifferentiated cells and this strong phenotype was observed in all (n=10) independent transformants, suggesting accumulation of monomeric MpARF2 strongly interferes with normal development. This finding supports the notion that, in addition to favouring MpARF1:MpARF2 stoichiometry, proteasomal degradation prevents the accumulation of monomeric MpARF2.

## Discussion

We used the simple, minimal auxin response system in Marchantia to map protein levels both quantitatively and qualitatively throughout gemma development. To this end, we generated a full set of genomic knock-in lines, fusing fluorescent proteins (in two colours) to the C-terminus of each component by homologous recombination. While such an effort would be realistic in Physcomitrium patens, given the facile gene targeting in that species^21^, but more complicated due to the larger gene families. We do not know of any other plant species that would currently allow such a holistic investigation of the auxin response system. Based on normal phenotypes, we believe these knock-ins to faithfully represent the native spatio-temporal accumulation profile of all nuclear auxin response proteins in Marchantia.

By following the localization of all components over time, we identified dynamics of MpTIR1 receptor, MpAux/IAA repressor, and IAA itself that suggests a progressive transition from a state with high auxin, low receptor, and low Aux/IAA to one with low auxin, high receptor, and higher Aux/IAA. Thus, the capacity for Aux/IAA-dependent auxin response dynamically changes during development. One interesting idea is that the gemmae cup itself provides a high-auxin environment, as proposed previously^22^ that effectively causes degradation of Aux/IAA proteins. The capacity to respond to auxin depends on both the TIR1-Aux/IAA “permissive” machinery and the ARF transcription factors^6,7^. While imaging native ARF levels, we found a surprising and profound degradation process that changes the ARF landscape during the first hours after germination. The permissive auxin conditions at early stage are countered by an ARF landscape that is relatively inhibitory. Over time, this ARF landscape shifts to a relatively activating state. The prediction would be that as time progresses, gemmae will become more sensitive to small changes in auxin concentrations, but this prediction remains to be tested.

We could connect ARF degradation to two phenomena: active control of A/B ARF stoichiometry and selective stabilization of dimeric ARFs. With regard to the former, even in the absence of any treatment, and in static observation, gemmae represent a rich landscape of MpARF1:MpARF2 stoichiometries. From first principles, and supported by mathematical modeling, these sites of varying stoichiometries should translate to areas with different auxin response outputs. Indeed, we see that the cells with the most “activating” stoichiometry are rhizoid initial cells, that are known to be highly sensitive to externally applied auxin^23–25^. Thus, the endogenous ARF accumulation patterns will likely translate to a corresponding map of auxin sensitivities. Unfortunately, these are hard to map at present due to the absence of a cellular-resolution reporter for mapping auxin response output. Given that manipulating the stoichiometry does prevent normal auxin response and development, we do expect that the maps – both in the gemma studied here and beyond – will be an exciting starting point to map sites of auxin action.

A key question is how changes in ARF stoichiometry are brought about. In principle, any gene/protein regulatory process can contribute to protein accumulation, and this stoichiometry therefore offers a central pivot point in controlling auxin output. While it remains to be seen what transcriptional inputs contribute to diverse ARF gene expression patterns, we do see that – given unequal starting levels – a relatively generic degradation rate can create large changes in MpARF1:MpARF2 stoichiometry. Identification of components in the degradation mechanism, as well as in (post)transcriptional control will help resolve the tuning mechanisms.

By mapping the degron motif in MpARF2, we made a surprising discovery: the region both necessary and sufficient for degradation is located at the interface between both DBD monomers in the protein dimer. While we cannot be certain that dimerization occludes the degron, it is extremely unlikely that this interface will be available for the proteolysis adaptor protein. Thus, this finding suggests a regulatory mode where degradation selectively targets MpARF2 monomers. We find that mutations that interfere with dimerization render the protein unstable, and this includes mutations in the C-terminal PB1 domain. This suggests that cooperative dimerization protects the protein from destruction and also helps rationalise the earlier findings that the MpARF1 PB1 domain is essential for function *in vivo* yet can be replaced by an entirely unrelated dimerization domain^11^. A mutant version of MpARF2 that can neither dimerize nor be degraded leads to extremely strong growth and developmental phenotypes, suggesting that indeed, monomers are detrimental to normal development. A key question is why this is the case. A plausible explanation is that the limited specificity of an ARF monomer for 4 conserved nucleotides (TGTC) allows binding of monomers across the genome (every 256 bases), while an inverted repeat of the same element would lead to a specificity that is orders of magnitude higher (every 65,536 bases). As such, a monomer could be a loose cannon in auxin-dependent gene regulation (Figure 6.E). While this interpretation is plausible, and to some degree testable, it is a mystery why this problem was not solved by evolving higher homotypic ARF dimerization affinity. Is the dynamic regulation of the monomer-dimer equilibrium important for auxin response, perhaps under environmental influences, or are there structural and functional constraints that we are yet to identify?

ARFs can bind to inverted (IR), everted (ER) and direct (DR) repeats of their binding site, with defined spacing^19,20,26,27^. The degron maps to the dimerization interface that is generated in unbound or IR-bound ARF’s. This conformation creates an ARF dimer where each monomer is a mirror image of the other. However, cooperative binding to DR or ER elements would likely require a different conformation, in which the degron is exposed. Therefore, selective MpARF2 monomer degradation may additionally act as a mechanism to favour binding to IR motifs.

Now that we have identified a degron motif for MpARF2, it will be interesting to see if the mechanisms underlying MpARF1 degradation work similarly. The region spanning the degron is highly conserved between MpARF1 and MpARF2, which makes it likely that the same binding partner mediates degradation of both proteins. If so, this would directly mean that the control of A/B ARF stoichiometry and monomer/dimer equilibrium are mechanistically connected. Identification of binding partners, such as E3 ubiquitin ligase subunits may help to address this question.

MpARF1 and MpARF2 are the sole representatives of the A-class and B-class ARFs in Marchantia, and both share ancestry with proto-A/B-ARFs in algal ancestors^10^. It is thus plausible that DBD-mediated degradation is a property inherited from the algal ancestor. This urges the question of how widespread this type of regulation is. Several of the 23 Arabidopsis ARFs (AtARF1, AtARF6, AtARF7, AtARF17 and AtARF19) have been reported to be proteasomally degraded^28,29,30^. Neither the degron nor the biological relevance of these degradation is known, but there is clearly a potential for this degradation mechanism to be intimately connected to auxin response.

Finally, with identifying ARF proteolysis, it now seems that all components in the NAP are subject to selective degradation. Aux/IAAs are degraded in an auxin-dependent manner^5^. Intriguingly, when not engaged in binding Aux/IAA proteins, Arabidopsis TIR1 protein is targeted from degradation^31^. Thus, auxin response is marked by profound proteasomal degradation, which may be a requirement for the system to rapidly adapt to changing internal and external conditions.

## Materials and methods

### Plant growth conditions

*Marchantia polymorpha* male Takaragaike-1 (Tak-1) and female Takaragaike-2 (Tak-2) plants were used as the wild-type variety. For vegetative propagation, plants were grown on ½ Gamborg B5 media plates in growth chambers with 40 μmol photons m^−2^ s^−1^ continuous white light at 22 °C. For sexual reproduction, plants were grown on 1% sucrose supplemented ½ Gamborg B5 media within hyrophonic boxes exposed to 40μmol photons m^−2^ s^−1^ continuous white fluorescent light for 1 month. Plants were then moved into 40 μmol photons m^−2^ s^−^ 1 continuous white light supplemented with 15μmol photons m^−2^ s^−1^ far-red light to induce antheridiophore and archegoniophore development. Plants were repeatedly crossed manually to ensure fertilization. Sporangia with mature spores were collected aseptically and used in spore transformation.

### Development of genomic knock-in translational fusions

Marchantia knock-in lines were developed to study native auxin response proteins at endogenous concentrations. A NOS terminator and a fluorescent marker gene (either mScarlet-I or mNeonGreen) were cloned sequentially at the HindIII restriction site of pJHY-TMp1 binary plasmid to create pJHY-mScarlet-I and pJHY-mNeonGreen vectors. After each cloning step, the HindIII site was regenerated by adding a HindIII site in the forward primer. This allowed subsequent cloning at the 5’ end of the previous insert in the same plasmid. Two 3.5kb genomic DNA fragments were amplified by PCR and used as homologous arms for recombination. The first genomic DNA fragment contained the 3.5kb sequence upstream of the stop codon of the gene of interest while the second fragment was composed of the 3.5kb sequence downstream of the stop codon. The first fragment was cloned at the HindIII site and the second fragment was cloned at the AscI site of pJHY-mScarlet-I and pJHY-mNeonGreen. This cloning strategy was used to create homologous recombination constructs for ARF1, ARF2, ARF3, TIR1, Aux/IAA and ncARF. Wild-type (Tak-1) spores were transformed by agrobacterium mediated transformation protocol described by Ishizaki et. al. (2008)^32^. Transformants were selected on 1\2 Gamborg B5 + 100mg/litre Cefotaxime medium with 10mg/L hygromycin selection. Genomic DNA PCR was used to isolate true knock-in lines.

### Auxin sensitivity and physiological analysis of knock-ins

Knock-in lines were tested for their wild-type like growth, physiology, and auxin sensitivity. Tak-1, Tak-2 and all knock-in lines were treated with either DMSO or 3μM NAA supplemented ½ B5 media and grown for 7 days. On 8^th^ day, plants (n=10 per genotype) were imaged with stereomicroscope to compare their physiological responses to auxin (Supplementary Figure 1).

### Microscope slide mount setup for time course imaging

A microscope slide mount was set up for live imaging of gemmae to precisely track a selected set of cells for temporal protein expression analysis. The mount consisted of a circular aluminium disc with a plastic inset fitted at the centre of the disc (Supplementary Figure 2). Melted ½ B5 media with or without desired treatments were poured into the cavity of the plastic inset and allowed to solidify. Gemmae were carefully placed on top of the solidified media and covered with a round coverslip. The bottom of the mount was sealed with parafilm to prevent any evaporation and media drying during time series imaging. Between two imaging time points in a time series experiment, the mounts were placed inverted in the growth chambers to keep the gemmae exposed to light and allow normal growth.

### Confocal live cell imaging

All live cell imaging was done on a Leica SP8X-SMD confocal microscope equipped with hybrid detectors and a pulsed (40MHz) white-light laser (WLL). mNeonGeen and mScarlet-I were excited with the 506 nm and 561 nm laser lines, respectively. The laser powers were set at 4% output to avoid bleaching of the fluorophores. Fluorescence was detected between 512-560nm (mNeonGreen) and 570 to 620nm (mScarlet-I) using hybrid detectors in photon counting mode. Z-stack images of 1.5μm were acquired using a 20X water immersion objective lens and time-gated detection to suppress chlorophyll autofluorescence. Images were processed using ImageJ software. Maximum-intensity projections of z-stack images were used to quantify total cellular fluorescence in each nucleus analysed, corrected for background fluorescence.

### RNA extraction, cDNA synthesis and qPCR

Total plant RNA was extracted from gemmae that were collected from gemma cups of 4-week-old Tak-1 and knock-in plants and subsequently incubated in liquid half-strength B5 medium for 0, 3.5, and 8 hours, before freezing in liquid nitrogen. RNA was extracted from ground tissue using the TRIzol reagent and Qiagen Plant mini kit. An on-column RNase-free DNase (Qiagen) treatment was performed before final elution. cDNA was synthesized from 1 μg total RNA using the iScript Reverse Transcriptase kit (Biorad). qPCR reactions were carried out 2x IQ SYBR green (Biorad) on a CFX384 Touch Real-Time PCR detection system (Bio-Rad). Data analysis was performed as described by Taylor et al 2019^33^. Housekeeping genes *MpEF1a* and *MpSAND* were used for transcript level normalization.

### Inducible ARF2 overexpression

For inducible ARF2 overexpression, the pEF1∷ARF2-GR lines were used^11^. Plants (n=10 per genotype) were treated with either DMSO or 1μM dexamethasone in B5 media and imaged after 3 days to look for rhizoid formation as an indicator of gemma germination. Dexamethasone treatment was used to induce the movement of ARF2-GR from cytosol to nucleus.

### Mathematical modelling parameters and assumptions

Please see Supplementary file 1

### ARF2 domain split analysis

To locate the degron within ARF2 protein structure, each domain of ARF2 was expressed separately in plants. In pJHY-TMp1 vector backbone, a ~1.1kb promoter of ARF2 was cloned with respective DBD or MR or PB1 domains or full length ARF CDS fused to an mNeonGreen fluorescent marker. Each CDS was fused to an N-terminal SV40 NLS sequence to ensure nuclear localization of the protein. All primers were designed with appropriate overlapping overhangs and HiFi DNA assembly master mix (NEB) was used for cloning. Constructs were transformed in wild-type spores and transformants were selected on hygromycin selection plates.

### ARF2 DBD fragment analysis

To narrow down the core degron region within ARF2 DBD, the DBD was divided into 8 fragments of 75aa each. Each fragment had a 25aa overlap with its preceding and succeeding fragments. All 8 fragments, fused to an N-terminal SV40NLS sequence, were cloned at the HindIII site of pJHY-mNeonGreen plasmid. A ~2kb ARF2 promoter was cloned upstream of each fragment. These constructs were used to generate 8 transgenic lines expressing different parts of the ARF2 protein fused with mNeonGreen marker.

### ARF2 dimerization mutant analysis

ARF2 dimerization mutant lines were created to test the stability of monomeric ARF2. Overlapping primers with mutated nucleotides were used to amplify ARF2 CDS and cloned at HindIII site of pJHY-mNeonGreen plasmid, using NEB HiFi cloning master mix. G286I (DBD dimerization mutation), K760S+OPCA (PB1 dimerization mutation), and G286I+K760S+OPCA double dimerization mutant ARF2 CDS were cloned in frame with mNeonGreen tag and under the pARF2 promoter. Each construct was transformed in WT spores to generate 3 different dimerization mutant variants. Gemmae of these plants were used to visualize mutant ARF2 expression and stability during dormancy and growth.

### ARF2 DBD degron swap with ARF3 DBD sequence

The 25aa overlapping region between fragment 6 and fragment 7 of ARF2 DBD were divided into 3 fragments of 12aa each. These 12aa fragments from ARF2 DBD were swapped with the aligned sequences from ARF3 DBD. Overlapping primers with ARF3 sequence at 5’ end, were used to amplify the ARF2 CDS and fragments were cloned into pJHY-mNeonGreen using NEB HiFi cloning master mix.

### Total Indole-3 -acetic acid (IAA) and oxidized IAA quantification

Total IAA was quantified to estimate the total cellular auxin levels during gemma growth. Gemmae were collected from 4-week-old Tak-1 plants. Tak-1 gamme were grown on liquid ½ B5 media and samples were collected after 0, 3.5, 8 and 24 hrs of growth. Samples were snap frozen in liquid Nitrogen and ground into fine power and weighed. For the extraction of indole-3-acetic acid (IAA) ~150 mg of snap-frozen plant material was used per sample. Tissue was ground to a fine powder at −80°C using 3-mm stainless steel beads at 50 Hz for 2*30 seconds in a TissueLyser LT (Qiagen, Germantown, USA). Ground samples were extracted with 1 mL of cold methanol containing [phenyl 13C6]-IAA (0.1 nmol/mL) as an internal standard in a 2-mL eppendorf tube as previously described^34^. Sample were filtered through a 0.45 μm Minisart SRP4 filter (Sartorius, Goettingen, Germany) and measured on the same day. IAA was measured on a Waters Xevo TQs tandem quadruple mass spectrometer.

## Acknowledgement

We are thankful to Iris Nieuwland for her help with the generation of MpARF3-mScarlet-I knock-in, Juriaan Rienstra for assisting in the ARF2 degron mapping in PyMol and Neri van Laar for performing the Bortezomib experiment on ARF knock-ins. We acknowledge NWO-ALW (Project number ALWOP.402) for funding this project.

---

**Supplementary Figure 1:**
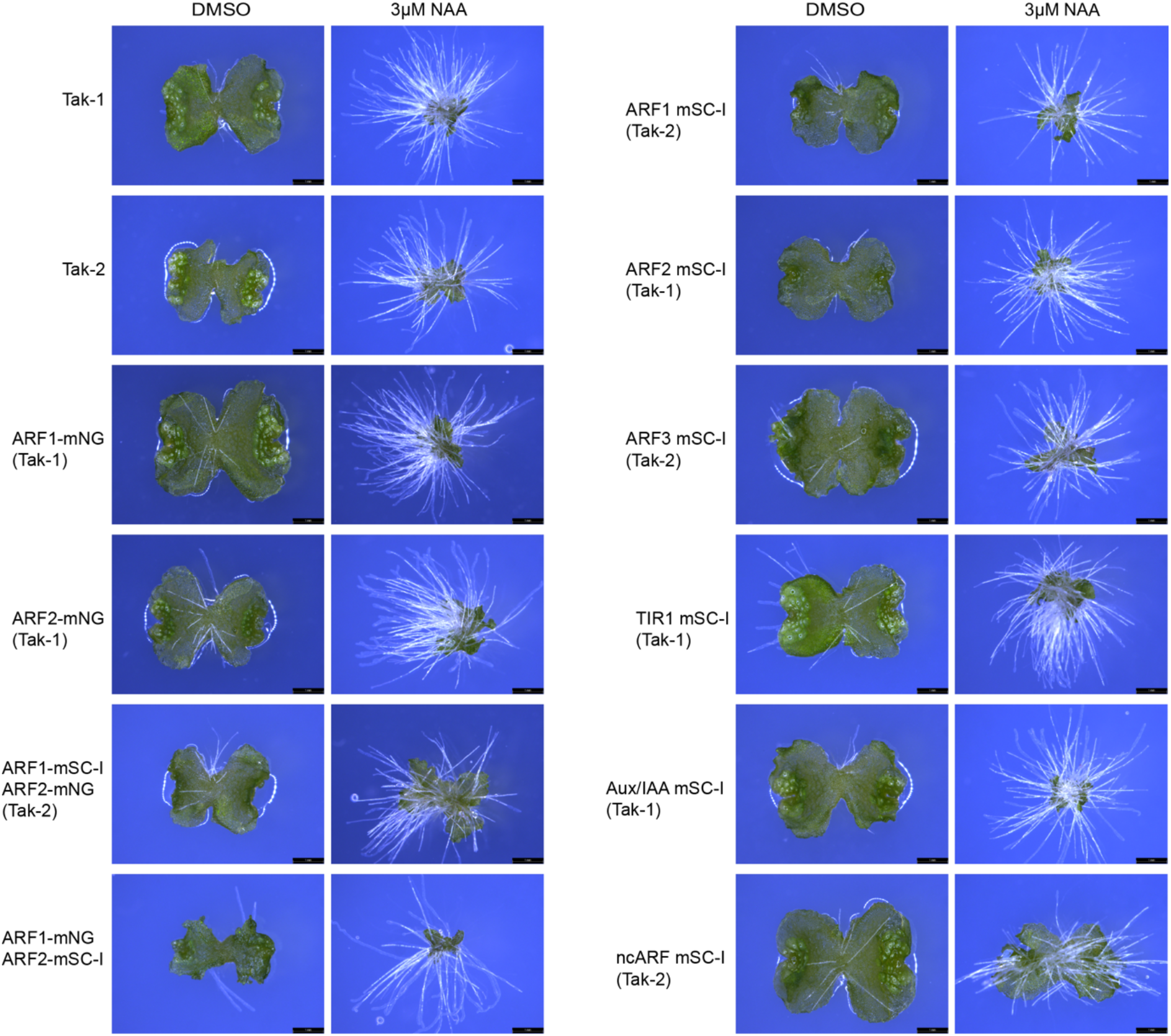
Auxin response assay on all genomic knock-in lines. All knock-ins were treated with DMSO or 3μM NAA and imaged after 1 week. All knock-ins show wild-type like auxin response with thallus growth inhibition and ectopic rhizoids formation.

**Supplementary Figure 2:**
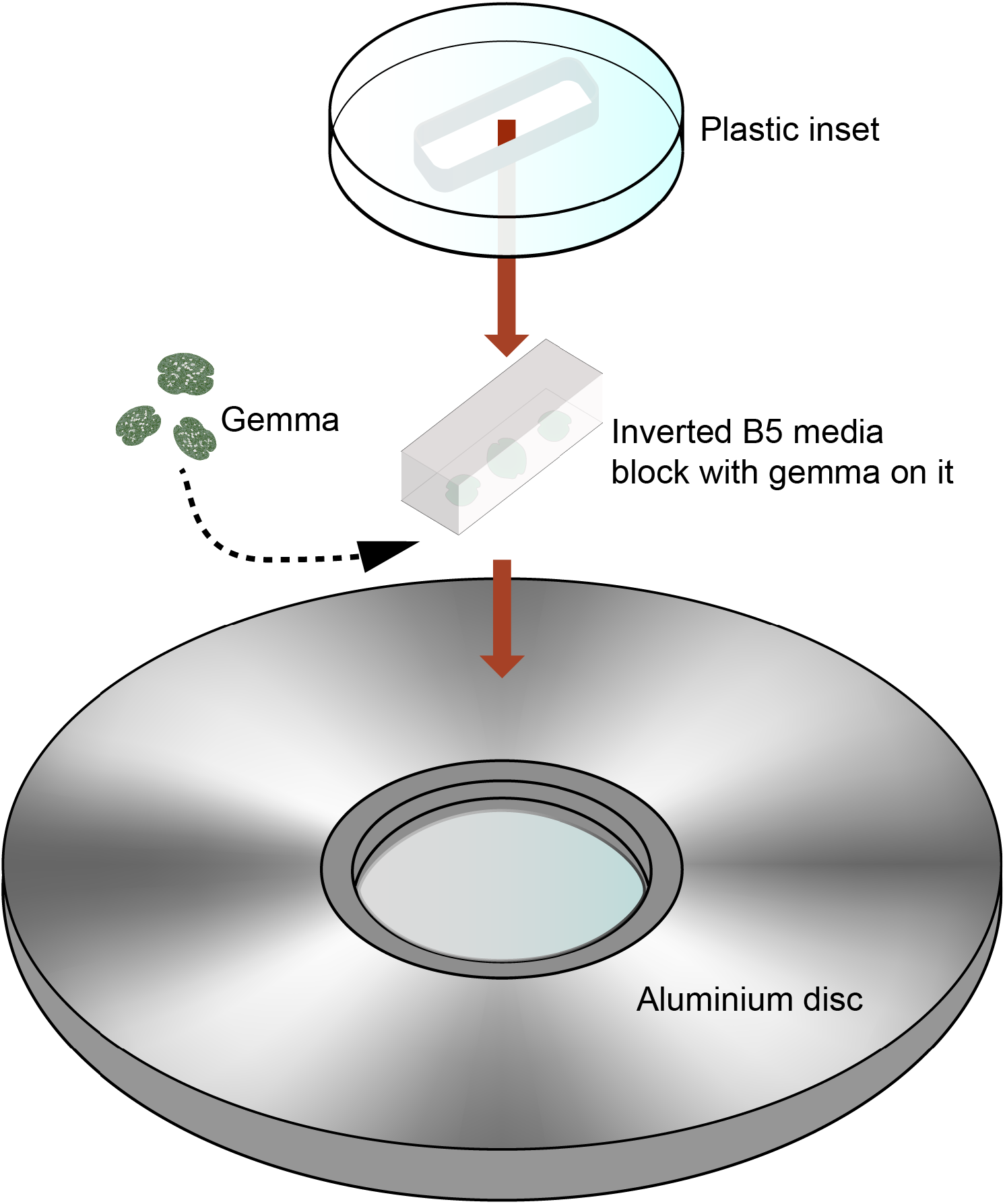
Basic design of the microscope slide mount, used for time course imaging. B5 media blocks are solidified inside the plastic inset and gemmae are places on top of the media. The media is supplemented with desired treatment or mock before casting in the inset. A round coverslip is gently placed on top of the gemmae. The plastic inset containing the gemma samples on the media block, is inverted and placed on the aluminium disc and tightened with a screw to prevent movement. Evaporation of water from the media block is prevented by sealing the reverse side of the block with parafilm.

**Supplementary Figure 3:**
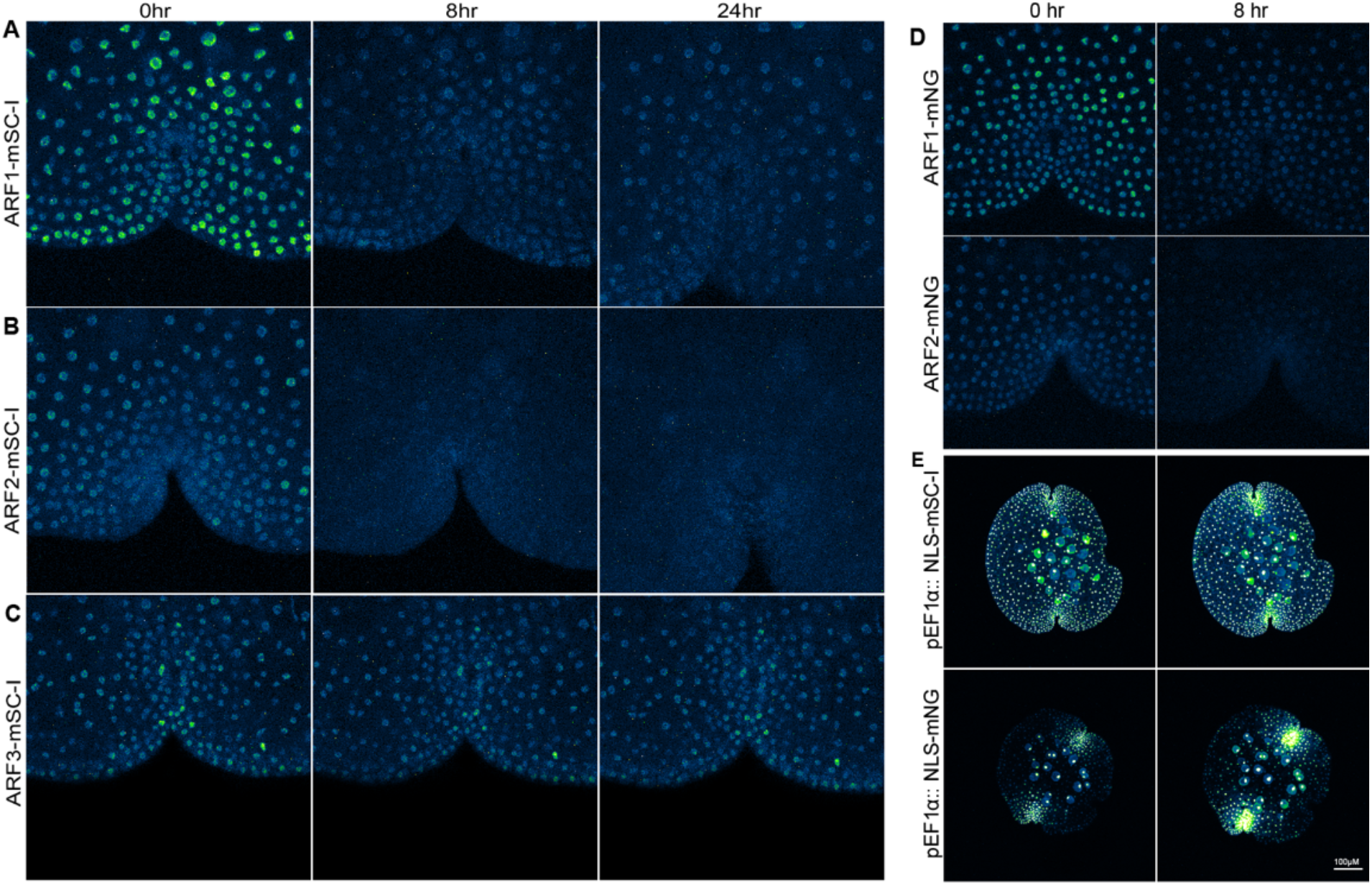
Time course imaging on ARF1, ARF2 and ARF3 -mScarlet-I (mSC-I) single knock-in lines. (A-B) ARF1 and ARF2 shows fluorescence decline after gemma germination. (D) Both mNG and mSC-I fusion variants show similar expression dynamics. (C) ARF3-mSC-I protein accumulation remains stable in the first 24 hours. (E) Non-fused fluorescent proteins are stably expressed at same developmental stages of gemmae, indicating specificity of fluorescence decline for ARF1 and ARF2.

**Supplementary Figure 4:**
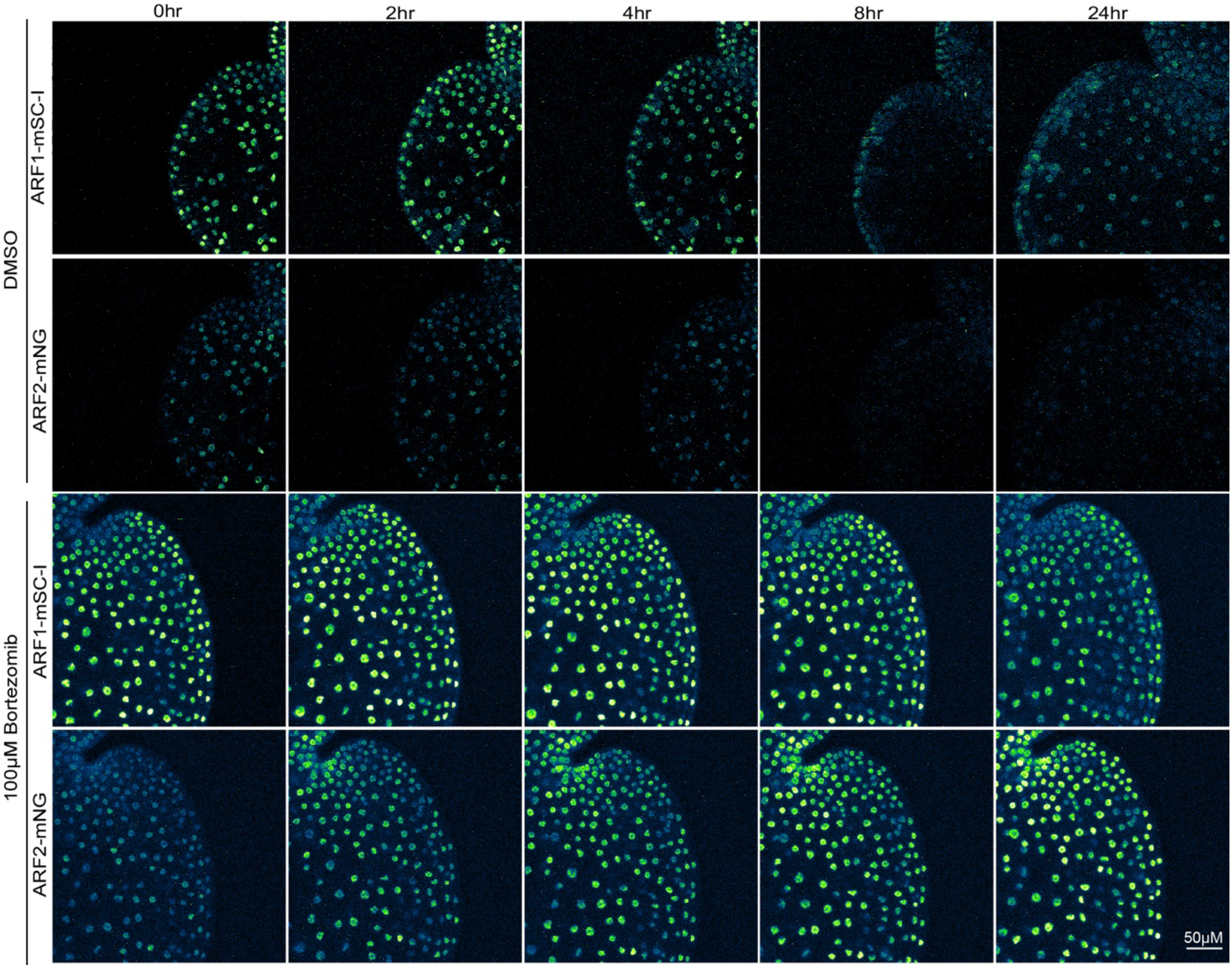
Treatment with an alternative proteasomal degradation inhibitor Bortezomib also blocks the degradation of ARF1-mSC-I and ARF2-mNG.

**Supplementary Figure 5:**
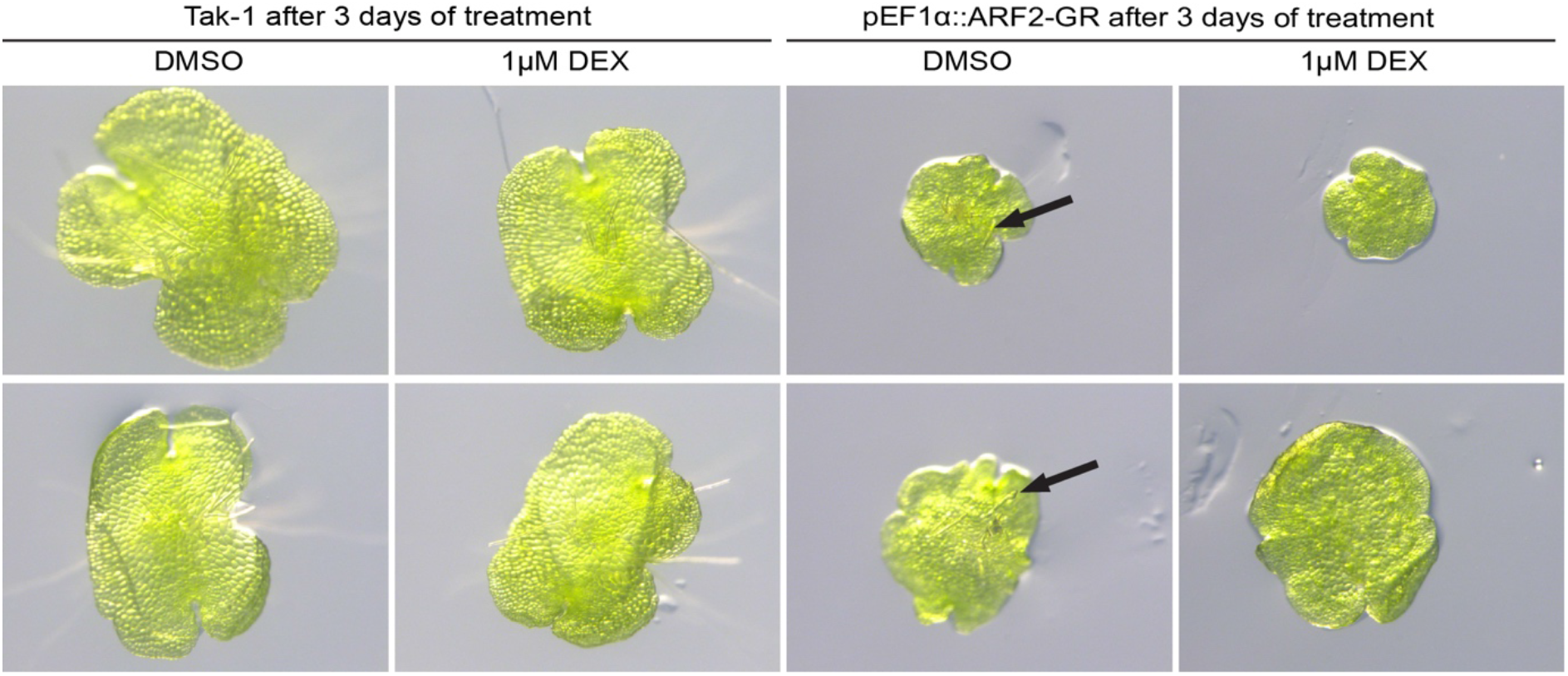
Inducible ARF2 overexpression leads to delayed gemma germination. pEF1∷ARF2-GR lines were treated with 1μM dexamethasone and plants were checked for rhizoid formation as a sign for germination. After 3 days of DMSO treatment, both wild-type Tak-1and pEF1∷ARF2-GR lines developed rhizoids. However, only Tak-1 developed rhizoids on dexamethasone treatment. No rhizoid was formed in induced ARF2-GR overexpression lines.

**Supplementary Figure 6:**
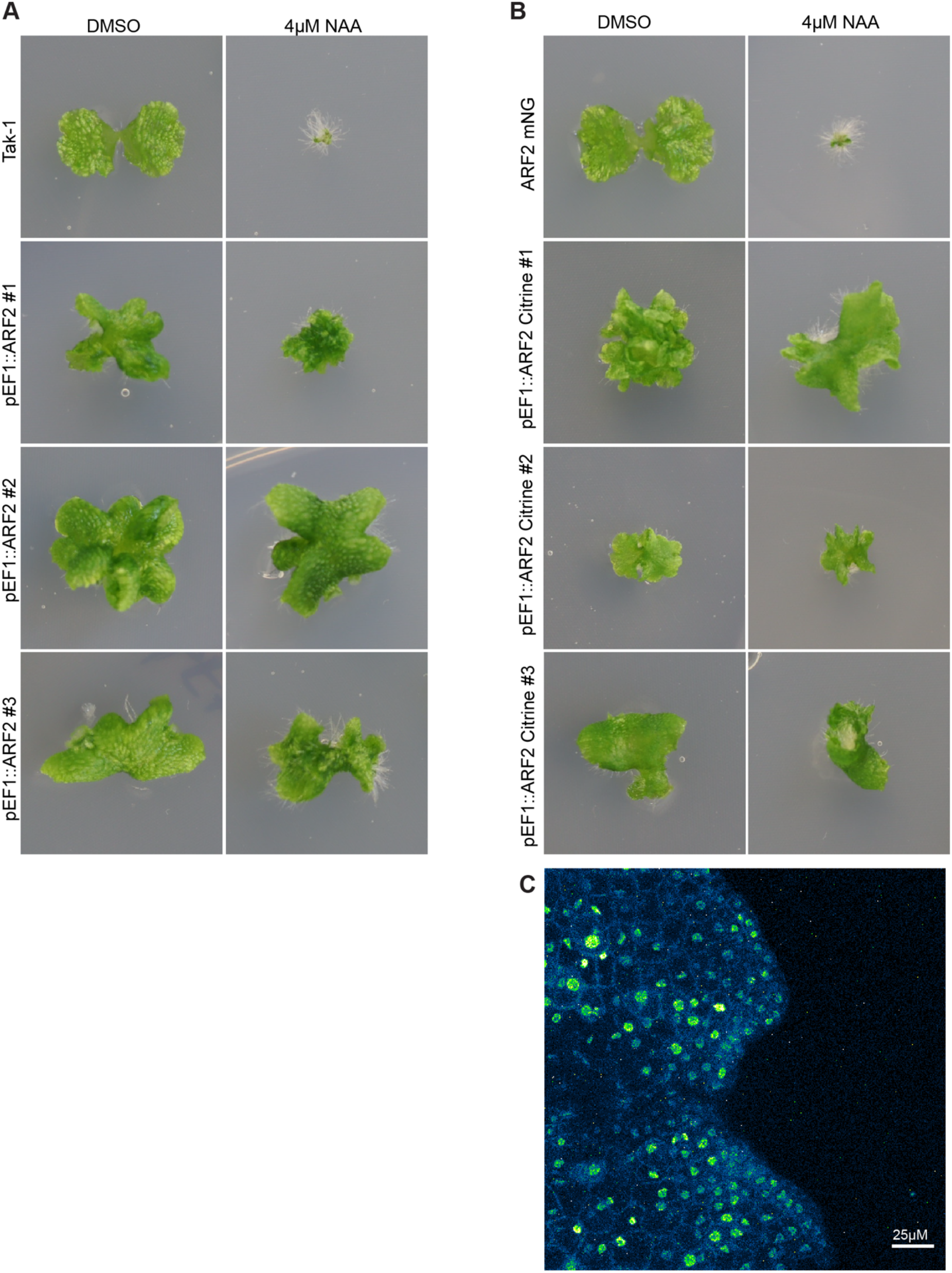
Constitutive overexpression of ARF2 or ARF2-Citrine leads to auxin insensitivity. (A) Three independent pEF1∷ARF2 lines showing auxin resistance in comparison with Tak-1 control. (B) Three independent pEF1∷ARF2-Citrine lines showing auxin insensitivity in comparison to ARF2-mNG knock-in. (C) Expression of ARF2-Citrine in pEF1∷ARF2-Citrine line#1 confirms that the auxin resistance phenotype is indeed due to the high expression of ARF2.

## Supplementary File 1

In this supplement the initial goal was to create an auxin signaling model for Marchantia to test out hypotheses. To do so we have altered an established auxin signaling model, the Farcot model^35^, to better represent Marchantia specifically.

The model we present is specific to modeling transcription rates relative to MpARF1 and MpARF2 concentrations within a gemma in the first 8 hours after dormancy. Time dependent simulations are run via Python3.8 utilizing primarily SciPy and NumPy packages. Specifically using the ODE solver scipy.intergrate.odeint.

The model takes the form of the following equations:

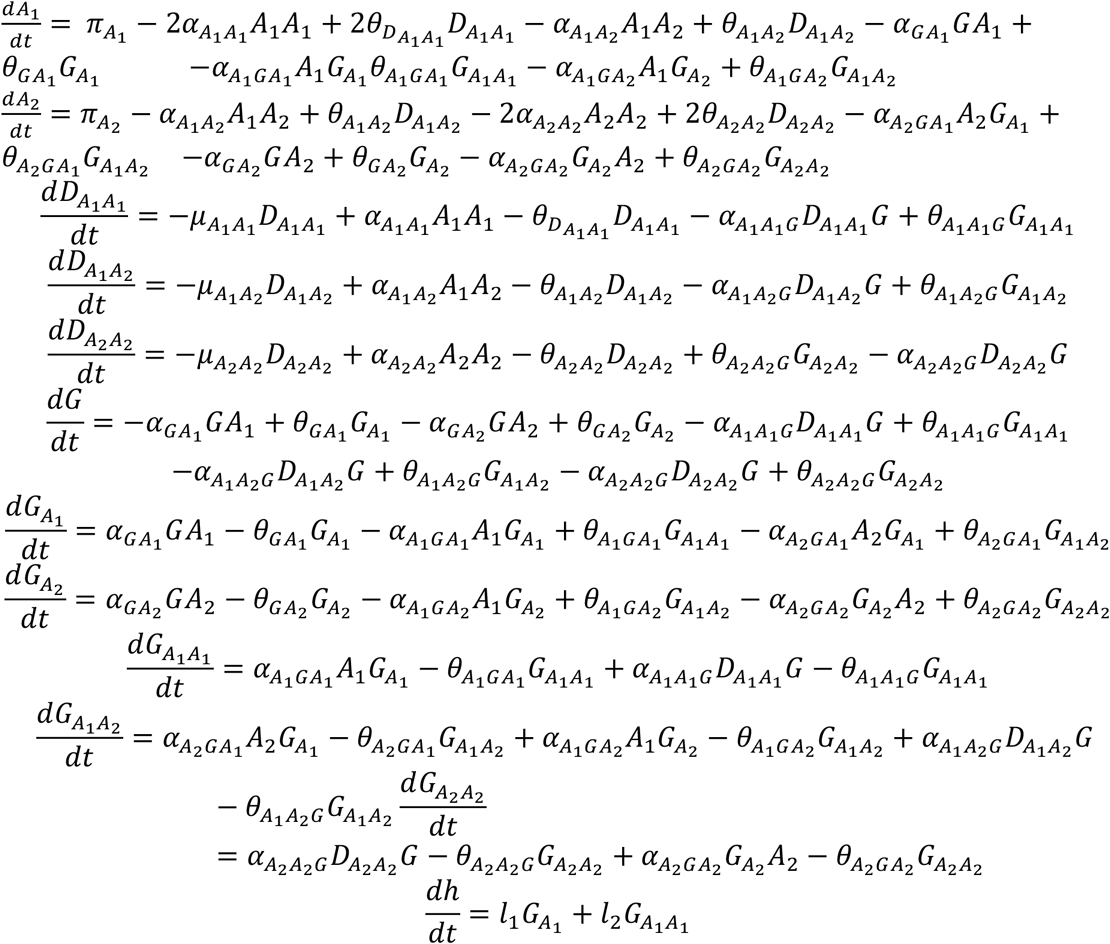

Where, *A*_1_ denotes the concentration of MpARF1, *A*_2_ denotes the concentration of MpARF2, *D*_*X,Y*_, *X*, *Y* ∈ {*A*_1_, *A*_2_} denotes the concentration of an X:Y dimer, 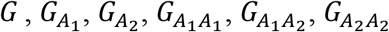 denote the proportion or the probabilities the promoter is free, bound with MpARF1, MpARF2, MpARF1:MpARF1 dimer, MpARF1:MpARF2 dimer or MpARF2:MpARF2 dimer respectively, *h* Denotes the concentration of mRNA being produced. The parameter values used are in the table below

| Variable | Value | Description |
| --- | --- | --- |
| $\alpha_{A_1A_1}$ | 1 | Rate of association of ARF1 with itself |
| $\alpha_{A_1A_2}$ | 1 | Rate of association of ARF1 with ARF2 |
| $\alpha_{A_2A_2}$ | 1 | Rate of association of ARF2 with itself |
| $\alpha_{A_1G}$ | $K_{on1}$ | Rate of association of an ARF1 and the promoter |
| $\alpha_{A_2G}$ | $K_{on2}$ | Rate of association of an ARF2 and the promoter |
| $\alpha_{A_1A_1G}$ | $2K_{on1}$ | Rate of association of an ARF1:ARF1 dimer and the promoter |
| $\alpha_{A_1A_2G}$ | $K_{on1} + K_{on2}$ | Rate of association of an ARF1:ARF2dimer and the promoter |
| $\alpha_{A_2A_2G}$ | $2K_{on2}$ | Rate of association of an ARF2:ARF2 dimer and the promoter |
| $\theta_{A_1A_1}$ | 1 | Rate of disassociation of ARF1 with itself |
| $\theta_{A_1A_2}$ | 1 | Rate of disassociation of ARF1 with ARF2 |
| $\theta_{A_2A_2}$ | 1 | Rate of disassociation of ARF2 with itself |
| $\theta_{A_1G}$ | $K_{off1}$ | Rate of disassociation of an ARF1 and the promoter |
| $\theta_{A_2G}$ | $K_{off2}$ | Rate of disassociation of an ARF2 and the promoter |
| $\theta_{A_1A_1G}$ | $\frac{1}{2}K_{off1}$ | Rate of disassociation of an ARF1:ARF1 dimer and the promoter |
| $\theta_{A_1A_2G}$ | $(K_{off1}^{-1} + K_{off2}^{-1})^{-1}$ | Rate of disassociation of an ARF1:ARF2dimer and the promoter |
| $\theta_{A_2A_2G}$ | $\frac{1}{2}K_{off2}$ | Rate of disassociation of an ARF2:ARF2 dimer and the promoter |
| $\mu_{A_1A_1}, \mu_{A_1A_2}$<br>or $\mu_{A_2A_2}$ | 0.05 | Rate of decay of ARF oligomers |
| $l_1$ | 1 | Rate of transcription caused by an ARF1 monomer |
| $l_2$ | 10 | Rate of transcription caused by an ARF1:ARF1 dimer |
| $\pi_{A_1}(t)$ | $\frac{dTotalARF1}{dt}(t) + \mu_{A_1A_2} + 2\mu_{A_1A_1}$ | Term that keeps total mpARF1 concentration matching experimental data |
| $\pi_{A_2}(t)$ | $\frac{dTotalARF2}{dt}(t) + \mu_{A_1A_2} + 2\mu_{A_2A_2}$ | Term that keeps total mpARF2 concentration matching experimental data |

## Model assumptions

In the model, decrease in total concentration of ARFs is assumed to be via the removal of ARF monomers. This is achieved via the 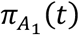 and 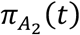 terms and is inspired by our understanding that the area of an ARF targeted for destruction is the DNA binding domain. We assume in this model that there is no effect on the rate of transcription caused by MpIAA due to its degradation in the early hours gemmae growth.

With respect to parameterization, we assume that the association and disassociation rate between ARFs is 1 for simplicity, and in absence of any quantitative experimental estimates. We assume ARF1 dimers are more effective in promoting transcription than ARF1 monomers and we assume ARF1 monomers and dimers containing ARF2 do not promote transcription. With respect to the association and disassociation of ARF molecules and the promoter we assume that dimers bind the ARF at a rate that is the summation of the rates of the constituent monomers. Furthermore, we assume that the rate of disassociation of dimers is the inverse of the sum of the inverses of the rates of dissociation of the constituent monomers. This in effect makes the rate of disassociation of dimers lower than that of either constituent monomer whilst considering how well either binds the promoter.

## Time series simulations

To produce time series simulations using the model for each individual time series dataset we set *TotalARF*1 and *TotalARF*2 equal to a piecewise linear interpolation of the time series data and used the parameters above. To compare the effect the differing Kds of MpARF1 and MpARF2 had on the system we varied *K_on1_ K_on2_ K_off1_ K_off2_* in 10 different combinations to represent either ARF having a greater Kd than the other whilst also altering the association and disassociation rates.

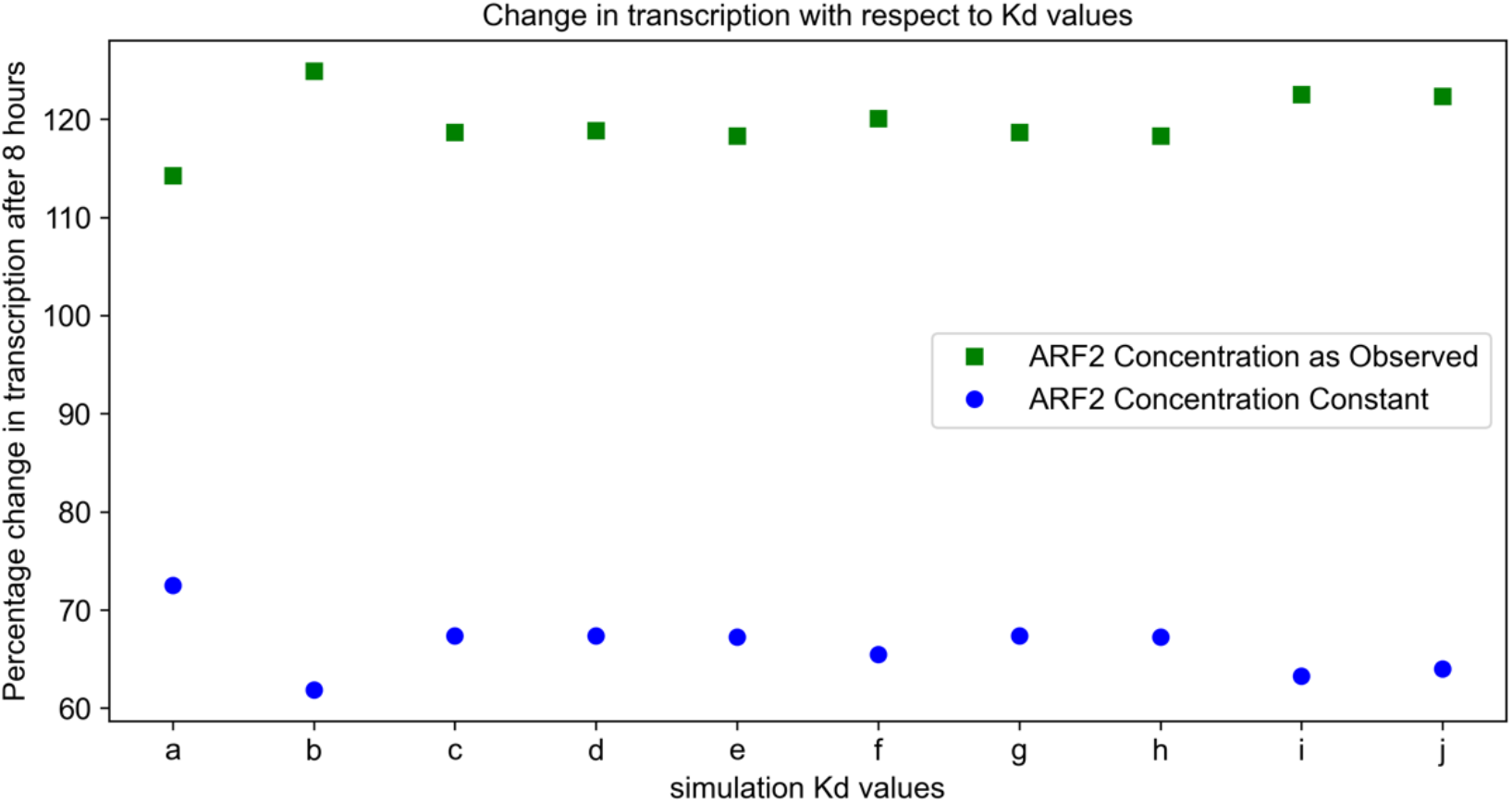
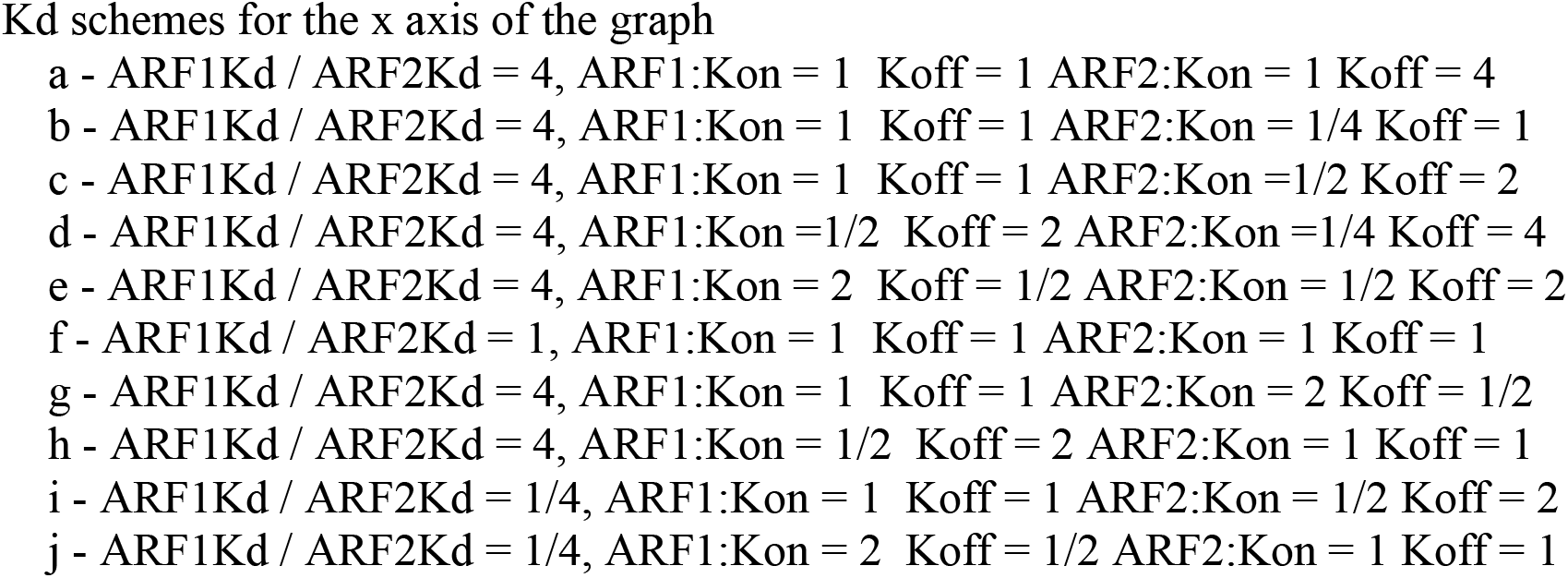

